# When Less is More: “Slicing” Sequencing Data Improves Read Decoding Accuracy and *De Novo* Assembly Quality

**DOI:** 10.1101/013425

**Authors:** Stefano Lonardi, Hamid Mirebrahim, Steve Wanamaker, Matthew Alpert, Gianfranco Ciardo, Denisa Duma, Timothy J. Close

## Abstract

Since the invention of DNA sequencing in the seventies, computational biologists have had to deal with the problem *de novo* genome assembly with limited (or insufficient) depth of sequencing. In this work, for the first time we investigate the opposite problem, that is, the challenge of dealing with excessive depth of sequencing. Specifically, we explore the effect of ultra-deep sequencing data in two domains: (i) the problem of decoding reads to BAC clones (in the context of the combinatorial pooling design proposed in [1]), and (ii) the problem of *de novo* assembly of BAC clones. Using real ultra-deep sequencing data, we show that when the depth of sequencing increases over a certain threshold, sequencing errors make these two problems harder and harder (instead of easier, as one would expect with error-free data), and as a consequence the quality of the solution degrades with more and more data. For the first problem, we propose an effective solution based on “divide and conquer”: we “slice” a large dataset into smaller samples of optimal size, decode each slice independently, then merge the results. Experimental results on over 15,000 barley BACs and over 4,000 cowpea BACs demonstrate a significant improvement in the quality of the decoding and the final assembly. For the second problem, we show for the first time that modern *de novo* assemblers cannot take advantage of ultra-deep sequencing data.

## Introduction

We have recently introduced in [1] a novel protocol for clone-by-clone *de-novo* genome sequencing that leverages recent advances in combinatorial pooling design (also known as *group testing*). In our sequencing protocol, subsets of non-redundant genome-tiling BACs are chosen to form intersecting pools, then groups of pools are sequenced on an Illumina sequencing instrument via low-multiplex (DNA barcoding). Sequenced reads can be assigned/decoded to specific BACs by relying on the combinatorial structure of the pooling design: since the identity of each BAC is encoded within the pooling pattern, the identity of each read is similarly encoded within the pattern of pools in which it occurs. Finally, BACs are assembled individually, simplifying the problem of resolving genome-wide repetitive sequences.

In [1] we reported preliminary assembly statistics on the performance of our protocol in four barley (*H. vulgare*) BAC sets (Hv3–Hv6). Further analysis on additional barley BAC sets and two genome-wide BAC sets for cowpea (*V. unguiculata*) revealed that the raw sequencing data for some datasets was of significantly lower quality (i.e., higher sequencing error rate) than others. We realized that our decoding strategy solely based on the software HASHFILTER [1] was insufficient to deal with the amount of noise in poor quality datasets. We attempted to (i) trim/clean the reads more aggressively or with different methods, (ii) identify low quality tiles on the flow cell and remove the corresponding reads (e.g., tiles on the “bottom middle swath”), (iii) identify positions in the reads possibly affected by sequencing “bubbles”, and (iv) post-process the reads using available error-correction software tools (e.g., QUAKE, REPTILE). Unfortunately, none of these steps accomplished a dramatic increase in the percentage of reads that could be assigned to BACs, indicating that the quality of the dataset did not improve very much. These attempts to improve the outcome led however, to a serendipitous discovery: we noticed that when HASHFILTER processed only a portion of the dataset, the proportion of assigned/decoded reads increased. This observation initially seemed counterintuitive: we expected that feeding less data into our algorithm meant that we had less information to work with, thus decrease the decoding performance. The explanation is that when data is corrupted, more (noisy) data is not better, but worse.

The study reported here directly addresses the observation that when dealing with large quantities of imperfect sequencing data, “less” can be “more”. More specifically we report (i) an extensive analysis of the tradeoff between the size of the datasets and the ability of decoding reads to individual BACs; (ii) a method based on “slicing” datasets that significantly improves the number of decoded reads and the quality of the resulting BAC assemblies; (iii) an analysis of BAC assembly quality as a function of the depth of sequencing, for both real and synthetic data. Our proposed algorithmic solution relies on a divide and conquer approach, as illustrated in Figure 1.

**Figure 1:**
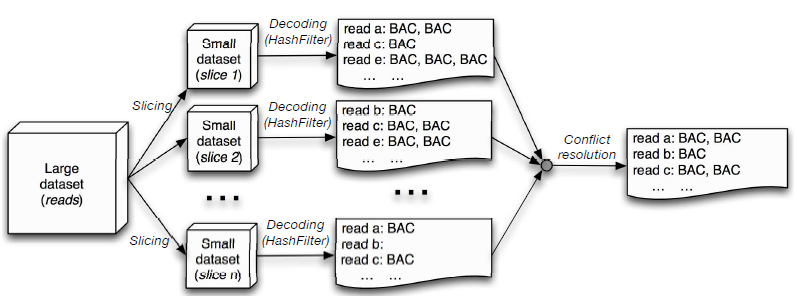
An illustration of the proposed strategy to improve read decoding: (1) a large dataset of reads to be decoded is “sliced” in *n* smaller datasets of optimal size, (2) each slice is decoded independently, and (3) read-to-BAC assignments for each slice are merged and conflicts are resolved

## Materials and Methods

We applied the combinatorial pooling scheme described in [1] to BAC clones for (i) a gene-enriched portion of the genome of *Hordeum vulgare L*. (barley), and (ii) the whole genome of *Vigna unguiculata* (cowpea). Briefly, in our sequencing protocol we (i) obtain a BAC library for the target organism; (ii) select gene-enriched BACs from the library (optional); (iii) fingerprint BACs and build a physical map; (iv) select a minimum tiling path (MTP) from the physical map; (v) pool the MTP BACs according to the shifted transversal design; (vi) sequence the DNA in each pool, trim/clean sequenced reads; (vii) assign reads to BACs (*deconvolution*); (viii) assemble reads BAC-by-BAC using a short-read assembler.

We should first note that a rough draft of the ≈ 5,300 Mb barley genome is now available [2]: our BAC sequencing work had contributed to that effort, but is distinct. In our work, we focused on the gene-enriched portion of the genome. We started with a 6.3x genome equivalent barley BAC library which contains 313,344 BACs with an average insert size of 106 kb [3]. About 84,000 gene-enriched BACs were identified and fingerprinted using high-information-content fingerprinting [4], [5]. From the fingerprinting data a physical map was produced [6], [7] and a MTP of about 15,000 clones was derived [8]. Seven sets of *N* = 2,197 clones were chosen to be pooled according to the shifted transversal design [9], which we called Hv3, Hv4, …, Hv9 (Hv1 and Hv2 were pilot experiments). An additional set of *N* = 1,053 clones (called Hv10) was pooled using the shifted transversal design with different pooling parameters (see below).

For barley sets Hv3, Hv4, …, Hv9, we chose parameters *P* = 13, *L* = 7 and 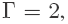, so that we could handle 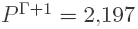 samples and make the scheme 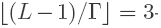 decodable. Each of the *L* = 7 layers consisted of *P* = 13 pools, for a total of 91 BAC pools. In this pooling design, each BAC is contained in *L* = 7 pools and each pool contains 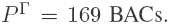 We call the set of *L* pools to which a BAC is assigned, the *BAC signature*. Any two BAC signatures can share at most 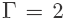 pools, and any triplet of BAC signatures can share at most 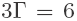 pools. For sets Hv3–Hv8, Vu1 and Vu2, we manually pooled 2,197 BACs thus exhausting all the “available” signature for the pooling design. However, for set Hv9 we only used 1,717 signatures. Set Hv10 was pooled using a different design: we chose pooling parameters *P* = 11, *L* = 7 and 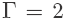 for a total of 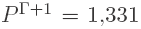 BAC signatures – however, we only used 1,053 signatures. BAC signatures that were available but not used in the pooling were called *ghosts*.

Cowpea’s genome size is estimated at 620 Mb and it is yet to be fully sequenced. For cowpea we started from a 17X depth of coverage BAC library containing about 60,000 BACs from the African breeding genotype IT97K-499-35 with an average insert size of 150 kb. Cowpea BACs were fingerprinted using high information content fingerprinting [4], [5], [10]. A physical map was produced from 43,717 fingerprinted BACs with a depth of 11x genome coverage [6], [11], and a minimal tiling path (MTP) comprised of 4,394 clones was derived [8]. The set of MTP clones was split in two sets of *N* = 2,197 BACs (called hereafter Vu1 and Vu2), each of which was pooled according to the shifted transversal design [9], with the same pooling parameters used for Hv3–Hv9.

To take advantage of the high throughput of sequencing of the Illumina HiSeq2000, 13–20 pools in each set were multiplexed on each lane, using custom multiplexing adapters. After the sequenced reads in each lane were demultiplexed, we obtained an average of 1,764 million reads in each set with a read length of about 92 bases and an insert size of 275 bases. Reads were quality-trimmed and cleaned of spurious sequencing adaptors, and then reads affected by *E. coli* contamination or BAC vector were discarded. The percentage of *E. coli* contamination averaged around 43%: as a consequence, the average number of usable reads after quality trimming and cleaning decreased to about 824 million, with an average high quality read length of about 89 bases. Table 1 reports the number of reads, number of bases, average read length, and *E. coli* contamination for each of the ten sets (Hv3, Hv4, Hv5, Hv6, Hv7, Hv8, Hv9, Hv10, Vu1 and Vu2).

**Table 1:** Basic statistics on the ten sequenced read datasets (seven for barley, two for cowpeas) analyzed in this manuscript

|  | *after demultiplexing* |  |  |  | *after demultiplexing/cleaning/trimming* |  |  |
| --- | --- | --- | --- | --- | --- | --- | --- |
|  | *reads (M)* | *bases (Mbp)* | *read len (bp)* | *% E.coli* | *reads (M)* | *bases (Mbp)* | *read len (bp)* |
| Hv3 | 2,476 | 227,773 | 92.00 | 41.12% | 1,240.2 | 110,056 | 88.74 |
| Hv4 | 1,363 | 125,273 | 91.91 | 39.36% | 713.4 | 63,384 | 88.85 |
| Hv5 | 1,142 | 105,089 | 92.00 | 51.11% | 505.1 | 45,088 | 89.27 |
| Hv6 | 1,133 | 104,239 | 92.00 | 65.96% | 282.4 | 24,970 | 88.42 |
| Hv7 | 2,288 | 210,535 | 92.00 | 46.11% | 928.9 | 82,503 | 88.82 |
| Hv8 | 1,802 | 165,803 | 92.00 | 44.04% | 730.8 | 64,651 | 88.46 |
| Hv9 | 1,596 | 146,816 | 92.00 | 40.66% | 736.2 | 65,697 | 89.24 |
| Hv10 | 971 | 89,370 | 92.00 | 20.95% | 748.2 | 67,600 | 90.36 |
| Vu1 | 2,475 | 227,696 | 92.00 | 36.66% | 1,208.1 | 108,666 | 89.95 |
| Vu2 | 2,402 | 221,006 | 92.00 | 43.12% | 1,144.6 | 103,026 | 90.01 |

The 91 pools (77 for Hv10) of trimmed reads for barley and cowpea were processed using our *k*-mer based algorithm called HASHFILTER which is fully described in [1]. Briefly, HASHFILTER builds a hash table of all distinct *k*-mers in the 91 (or 77) pools of reads, and records for each *k*-mer the set of pools where it occurs. Then it processes each read individually: (i) a read is decomposed in its constitutive *k*-mers; (ii) the set of pools of each *k*-mer is fetched from the hash table, and matched against the BAC signatures (allowing for a small number of missing/extra pools); (ii) *k*-mer signatures that match a valid BAC signature are combined to produce the final BAC assignment for the read. Recall that since our pooling is 3-decodable, we can assign each read to up to three BACs.

For some of the datasets, the percentage of reads decoded using this procedure was very low (as low as 22%). The investigation to identify the cause of the problem and how to deal with it, is the main contribution of this manuscript. Several analyses on the sequencing data suggested that the quality of sequencing from flow cell to flow cell (and lane to lane) was highly variable. To complicate the matter further, somewhere in the middle of the sequencing of our samples, the sequencing chemistry for the Illumina HiSeq 2000 was switched, which we believe contributed to lower quality (but increased the quantity) of the sequencing data.

Our first objective was to get rid of “bad” sequencing data that hampered the decoding step and the assembly downstream. Unfortunately, without a reference genome (both barley and cowpea genome are still not fully sequenced/assembled) this cleaning step was not easy. While HASHFILTER can tolerate some level of noise, it appeared that the full datasets contained too much of it – with the consequence that only a small fraction of the reads would be decoded. To our initial surprise, running HASHFILTER on a fraction of the reads yielded higher decoding percentages, which suggested the idea to “slice” the data.

The next question was to study the dependency between the size of the dataset on the performance of the decoding algorithm. To this end, we took samples of the original 91 (or 77) set of reads in sizes of 0.5M, 1M, 2M, 3M, 4M and 5M reads (details on the sampling method can be found in the next section) and computed the percentage of reads decoded by HASHFILTER on these samples of increasing sizes. Figure 2-A shows the percentages of decoded reads for sets Hv3, Hv4, Hv5, Hv6 and Hv7; Figure 2-B is for Hv8, Hv9, Hv10, Vu1 and Vu2. The x-axis is the number of reads per pool (in millions) given in input to HASHFILTER (*k* = 26). The rightmost point on these graphs corresponds to the full dataset. The datasets used to generate Figure 2 are available as Supplemental Dataset 1.

**Figure 2:**
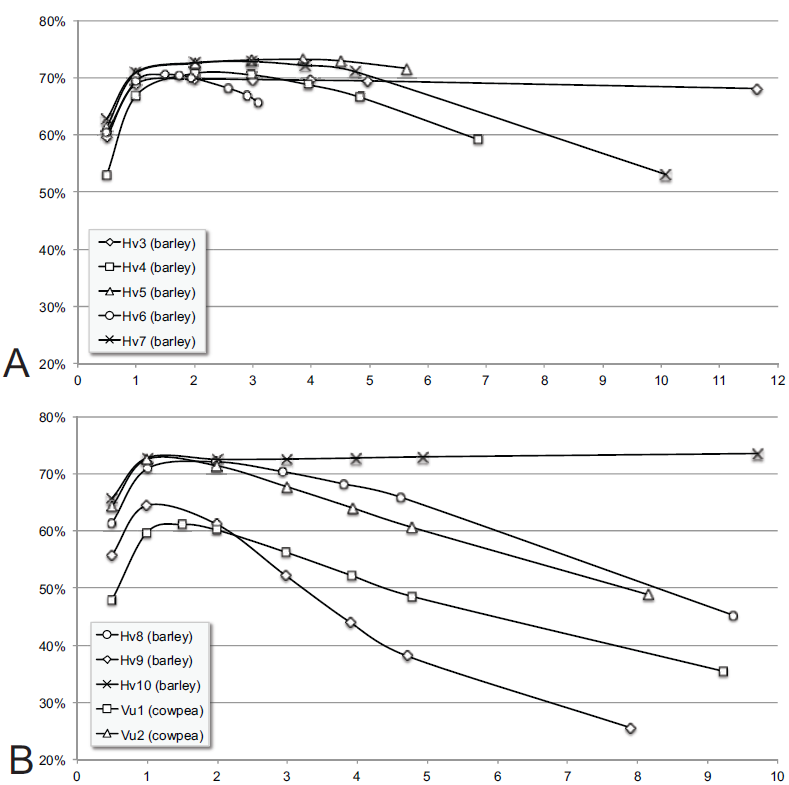
The percentage of reads decoded by HASHFILTER (*k* = 26) on dataset (A) Hv3, Hv4, Hv5, Hv6 and Hv7 (B) Hv8, Hv9, Hv10, Vu1, and Vu2 as a function of the number of reads given in input (X: number of million of reads sampled in each dataset)

Several observations on Figure 2 are in order. First, observe that when the number of reads per pool is too small (0.5-1M) the percentage of reads decoded by HASHFILTER is low. Similarly, when the number of reads per pool is large, the percentage of reads decoded by HASHFILTER can be low for some datasets. We believe that when the input size is small, there is not enough information in the hash table of *k*-mers to accurately decode the reads. However, when the input size is large, sequencing errors in the data introduce spurious *k*-mers in the hash table, which has the effect of deteriorating HASHFILTER’s decoding performance. Observe that almost all these curves reach a maximum in the range 1M–3M reads. For datasets whose “optimal number” of reads is low, we can speculate the amount of sequencing error to be higher. Also observe the large variability among these ten datasets. At one extreme, graphs for Hv3, Hv10 and Hv5 are very “flat” indicating low sequencing errors; at the other extreme, graphs for Vu1 and Vu2 degrade very quickly after the peak, indicating poorer data quality.

We also carried out a simulation study using synthetic reads generated from the rice genome (Oryza sativa). For this simulation we started from an MTP containing 3,827 BACs with an average length of about 150 kb, which spanned 91% of the rice genome (which is about 390 Mb). We pooled in silico a subset of 2,197 BACs from the set above according to the shifted transversal design (see [1] for details). We generated 2M synthetic reads for each of the 91 resulting rice BAC pools. Reads were 104 bases long with 1% sequencing error distributed uniformly along the read. A total of 208 Mbp gave an expected 56x coverage for each BAC. We ran HASHFILTER on the read datasets in slices of 0.25M, 0.5M, 1M, 1.5M and 2M (full dataset). The percentage of decoded reads (see Supplemental Text, Figure S1) peaks at 1.5M, and mirrors the observations made on real data. Even for synthetic reads, more data does not necessarily imply improved decoding performance.

## Improved Decoding Algorithm

Our improved decoding algorithm first executes HASHFILTER on progressively larger samples of the dataset (e.g., 0.5M, 1M, 2M, 3M, 4M, 5M and full dataset) for a given value of *k*. Our sampling algorithm selects reads uniformly at random along the input file: simply taking a prefix of the dataset is probably not a good idea because reads in the file are organized according to their spatial organization on the flowcell, possibly introducing biases.

When the sample size is greater than the pool size, the entire pool is used for decoding. Otherwise, reads in pools larger than the sample size are uniform sampled in order to meet the sample size constraint. As a result of this process, the size of each pool in a “slice” will be at most the sample size, but some of the pools will be smaller. The objective is to find the sample size that maximizes the number of reads decoded by HASHFILTER.

We observed that the optimal value of the sample size is somewhat independent from *k* as long as it is chosen “reasonably large”, say *k* > 20 for large eukaryotic genomes. Figure 3 illustrates that running HASHFILTER with *k* = 20, 23, 26, 29 gives rise to parallel curves. In fact, one can save time by running HASHFILTER with smaller values of *k* in order to find the optimal data size.

**Figure 3:**
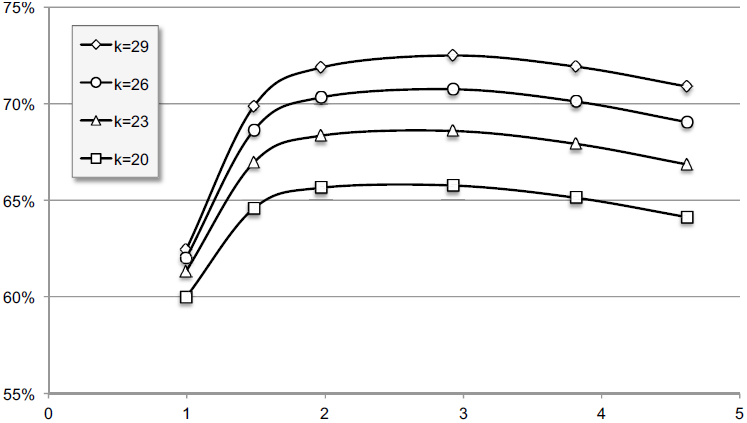
The percentage of reads decoded by HASHFILTER on one of the barley datasets for several choices of the *k*-mer size (X: number of million of reads sampled per pool)

Once the optimal sample size *N* is determined, the algorithm finds the size *M* of the largest pool to calibrate D datasets (hereafter called slices) each one of which has at most *N* reads per pool. For instance, if the optimal slice size is *n* = 2M reads, and the largest pool has *M* = 10M reads, the algorithm will create *D* = 5 slices: each one will be composed of 91 pools, each of which has at most 2*M* reads. Observe that the number of reads in each pool can vary significantly. For instance in Hv3, the largest pool has almost 23M reads, and the smallest has about 3M reads. Smaller pools will contribute their reads to multiple datasets. For instance, if there is a pool of size 2M in the same example above, these reads will appear in all five slices. In general, if *m* ≤ *n*, the entire pool will be used in each slice.

Then, the algorithms runs HASHFILTER *D* times, once on each of the D slices – which involves creating *D* individual hash tables. For this step, we recommend using the largest possible value of *k*, (*k* = 32), because the percentage of decoded reads for a given input size increases with *k* (see Figure 3). Then, the algorithm merges the D independent HASHFILTER’s outputs. If a read is decoded in only one slice, it will be simply copied in the output. If a read is decoded multiple times in different slices, the independent decodings may not agree, so a conflict resolution step is necessary. In our running example, reads in the small 2M-reads pool will be decoded five times: it is possible that HASHFILTER will assigned a read to five different BAC sets. In order to identify reads decoded multiple times, our algorithm first concatenates the *D* text outputs of HASHFILTER, then sorts the reads by ID, so that reads with the same ID are consecutive. We call a set of reads (single or paired-end) with the same ID, a read *group*. When a group contains more than one read, our algorithm computes the most likely assignment according to the following (conservative) rules. These rules are checked in order, the first one that applies is used, and subsequent rules are not considered.

1. if a group is composed of paired-end or single-end reads in which one or more BAC assignments has a 75%-majority, i.e., one or more BACs is associated with at least 75% of the reads in the group, then all the reads in the group are assigned to those majority BAC(s);
2. if a group is composed of at least two paired-end reads for which either the left or the right read is associated with one or more BACs with 50%-majority, then all the reads in the group are assigned to those BAC(s);
3. if a group is composed of paired-end reads for which either the left or the right read is not assigned, but at least 75% the assigned read agree on one or more BAC(s), then non-assigned mates are also assigned the same BAC(s);
4. if a group is composed of paired-end reads for which the BAC assignment for left and the right read disagree, then all BAC corresponding assignments are rejected.

Once all the decoded reads are assigned to 1–3 BACs using the procedure above, VELVET [12] is executed to assemble each BAC individually. As was done in [1], we evaluated the assembly for several choices of VELVET’s L-mer (hash) size (25–79, step of 6), but have reported only the assembly that maximizes the n50 (n50 indicates the minimum length of all contigs/scaffolds that together account for at least 50% of the genome). This is an arbitrary choice that does not guarantee the best overall assembly. In fact, based on our experience, when the sequencing depth is very high (say, over 500x) picking the assembly with the highest n50 has the effect of selecting “bloated” assemblies, i.e., assemblies whose sum of all contig lengths far exceed the target. In the next section we analyze the effect of excessive depth of sequencing on the assembly quality.

## Results

We employed several metrics to evaluate the improvement in read decoding and assembly enabled by the slicing algorithm. For one of the barley sets (Hv10) we executed HASHFILTER using several choices of *k* (*k* = 20, 23, 26, 29, 32) on the full 748M reads dataset (i.e., with no slicing) as well as with *k* = 32 using the slicing algorithm described above. The first five rows of Table 2 summarize the decoding results. First, observe that as we increase *k*, the number of decoded reads increases monotonically. However, if one fixes *k* (in this case *k* = 32, which is the maximum allowed by HASHFILTER), slicing Hv10 in 4 slices of ≈4M reads increases significantly the number of decoded reads (84.60% compared to 77.19%) available for assembly. Analysis of the number of assignments to ghost BACs also shows significant improvement in the decoding accuracy when using slicing: 0.000086% of the reads are assigned to unused BAC signatures compared to 0.000305%–0.001351% when HASHFILTER is used on the full dataset. We carried out a similar analysis on Hv9: when the full dataset was processed with HASHFILTER (*k* = 26), the number of reads assigned to ghost BACs was very high, ≈ 1.9M reads out of 196M (0.9653%). When the optimal slicing is used (*k* = 32), only 19,140 reads out of 516M are assigned to ghost BACs (0.0037%). Also, observe in Table 2 how the improved decoding affects the quality of the assembly for Hv10. When comparing no slicing to slicing-based decoding, the average n50 jumps from 12,260 bp to 42,819 bp (both for *k* = 32) and the number of reads used by VELVET in the assembly increases from 86.7% to 90.7%.

**Table 2:**
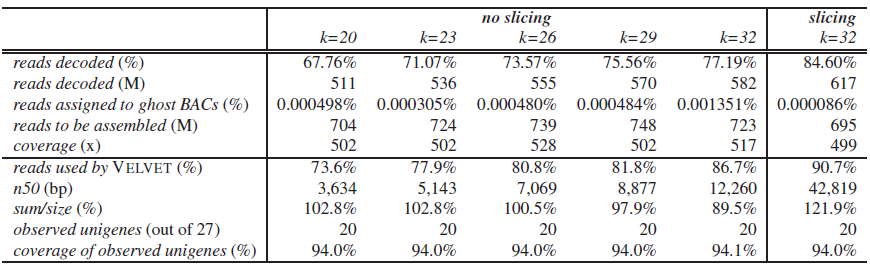
Decoding and assembly statistics for the Hv10 barley set for several choices of *k* on the full dataset, and for the improved slicing algorithm

|  | *no slicing* |  |  |  |  | *slicing* |
| --- | --- | --- | --- | --- | --- | --- |
| | $k=20$ | $k=23$ | $k=26$ | $k=29$ | $k=32$ | $k=32$ |
| *reads decoded (%)* | 67.76% | 71.07% | 73.57% | 75.56% | 77.19% | 84.60% |
| *reads decoded (M)* | 511 | 536 | 555 | 570 | 582 | 617 |
| *reads assigned to ghost BACs (%)* | 0.000498% | 0.000305% | 0.000480% | 0.000484% | 0.001351% | 0.000086% |
| *reads to be assembled (M)* | 704 | 724 | 739 | 748 | 723 | 695 |
| *coverage (x)* | 502 | 502 | 528 | 502 | 517 | 499 |
| *reads used by VELVET (%)* | 73.6% | 77.9% | 80.8% | 81.8% | 86.7% | 90.7% |
| *n50 (bp)* | 3,634 | 5,143 | 7,069 | 8,877 | 12,260 | 42,819 |
| *sum/size (%)* | 102.8% | 102.8% | 100.5% | 97.9% | 89.5% | 121.9% |
| *observed unigenes (out of 27)* | 20 | 20 | 20 | 20 | 20 | 20 |
| *coverage of observed unigenes (%)* | 94.0% | 94.0% | 94.0% | 94.0% | 94.1% | 94.0% |

For Hv10, we also measured the number of decoded reads that map (with 0,1,2 and 3 mismatches) to the assembly of a subset of 22 BACs that are available from [2]. Table 3 reports the average percentage of decoded reads (either from the full dataset or from the optimal slicing) that BOWTIE can map to the 454-based assemblies [2]. Observe how the slicing step improves by 6-7% the number of reads mapped to the corresponding BAC assembly, suggesting a similar improvement in decoding accuracy. Similar improvements in decoding accuracy was observed on the other datasets (data not shown).

**Table 3:** The percentage of reads that can be mapped (with BOWTIE with 0,1,2 and 3 mismatches) to the assembly of 26 BACs in Hv10 that are available from [2]

|  | *no slicing*<br>*k=32* | *slicing*<br>*k=32* |
| --- | --- | --- |
| *0 mismatches* | 75.2% | 82.4% |
| *1 mismatch* | 78.7% | 85.9% |
| *2 mismatches* | 80.5% | 87.4% |
| *3 mismatches* | 82.3% | 88.7% |

On Hv8, we investigated the effect of the slice size on the decoding and assembly statistics: earlier we claimed that the optimal size corresponds to the peak of the graphs in Figure 2. For instance, notice that the peak for Hv8 is ≈2M reads. We decoded and assembled reads using slicing sizes of 2M reads as well as (non-optimal) slice size of 3M reads. The experimental results are shown in Table 4. Observe that the decoding with 3M does not achieve the same decoding accuracy or assembly quality of the slicing with 2M, but again both are significantly better than without slicing. Again, notice in Table 4 how improving the read decoding affects the quality of the assembly. The average n50 increases from 4,126 bp (*k* = 26, no slicing) to 34,262 bp (*k* = 32, optimal slicing) and the number of reads used by VELVET in the assembly increases from 55.6% to 91.2%, respectively. For Hv8, 207 genes were known to belong to a specific BAC clone [1]: the assembly using slicing-based coding recovered at least 50% of the sequence of 187-190 of them, compared to 178 using no slicing.

**Table 4:** Decoding and assembly statistics for Hv8: comparing no slicing and slicing with two different slice sizes (2M reads is optimal according to the peak in Figure 2)

|  | *no slicing* | *slicing* |  |
| --- | --- | --- | --- |
|  | *k=26* | *k=32, 3M* | *k=32, 2M* |
| *reads decoded (%)* | 31.68% | 78.98% | 82.74% |
| *reads decoded (M)* | 270 | 539 | 600 |
| *reads to be assembled (M)* | 289 | 591 | 669 |
| *coverage (x)* | 94 | 197 | 223 |
| *reads used by VELVET (%)* | 69.0% | 92.6% | 91.6% |
| *n50 (bp)* | 4,126 | 31,226 | 34,262 |
| *sum/size (%)* | 55.6% | 97.0% | 102.0% |
| *observed unigenes (out of 207)* | 178 | 190 | 187 |
| *coverage of observed unigenes (%)* | 86.0% | 91.1% | 91.2% |

Finally, we compared the performance of our slicing method against the experimental results in [1], which were obtained by running HASHFILTER with no data slicing (*k* = 26). The basic decoding and assembly statistics when no slicing is used are reported in Table 5. First, observe the large variability of results among the ten sets. While the average number of decoded reads for *k* = 26 is ≈ 460M, there are sets which have less than half that amount (Hv6 and Hv9) and sets have more than twice the average (e.g., Hv3). As a consequence, the average fold-coverage ranges from 72x (Hv6) to 528x (Hv10). In general, the assembly statistics (without slicing-based decoding) are not very satisfactory: the n50 ranges from 2,630 bp (Hv9) to 8,190 bp (Hv3); the percentage of reads used by VELVET ranges from 66.0% (Hv9) to 85.9% (Hv3 and Hv4); the percentage of known genes covered at least 50% of their length by the assemblies ranged from 66% (Hv4) to 97% (Hv3).

**Table 5:** Decoding and assembly statistics for the ten datasets using *k* = 26 on the full dataset (no slicing)

|  | *decoding* |  | *assembly* |  |  |  |
| --- | --- | --- | --- | --- | --- | --- |
|  | *reads* (M) | *coverage* (x) | *n50* (bp) | *reads used* (%) | *sum/size* (%) | *observed/expected genes* (%) |
| Hv3 | 1,099.0 | 431.0 | 8,190 | 85.9% | 96.7% | 1,433/1,471 (97.42%) |
| Hv4 | 393.2 | 135.5 | 5,718 | 85.9% | 85.9% | 312/473 (65.96%) |
| Hv5 | 483.0 | 158.9 | 8,048 | 84.5% | 93.2% | 194/226 (85.84%) |
| Hv6 | 218.0 | 72.0 | 6,032 | 83.2% | 79.9% | 208/244 (85.25%) |
| Hv7 | 330.0 | 110.0 | 5,352 | 75.9% | 63.7% | 201/228 (88.16%) |
| Hv8 | 289.0 | 94.2 | 4,126 | 69.0% | 55.6% | 178/207 (85.99%) |
| Hv9 | 208.0 | 95.8 | 2,630 | 66.0% | 38.5% | 262/361 (72.58%) |
| Hv10 | 739.0 | 528.0 | 7,069 | 80.8% | 100.5% | 20/27 (74.07%) |
| Vu1 | 369.0 | 88.5 | 4,150 | 67.6% | 49.5% | 461/612 (75.33%) |
| Vu2 | 448.0 | 126.0 | 5,670 | 75.1% | 56.1% | 406/503 (80.72%) |
| *Average* | 457.6 | 184.0 | 5,699 | 77.4% | 72.0% | (81.13%) |

When we decoded the same ten datasets using the optimal slice size (using this time *k* = 32) the assemblies improved dramatically. The decoding and assembly statistics are summarized in Table 6: note that each set has its optimal size and the corresponding number of slices. First observe how the number of decoded reads increased significantly for most datasets (e.g., 330M to 785M for Hv7, 289M to 669M for Hv8, 209M to 516M for Hv9, 369M to 907M for Vu1 and 448M to 695M for Vu2). Only for two datasets the number of decoded reads decreased slightly (by 12M reads in Hv5, and by 44M in Hv10). For all the datasets, the average n50 increased dramatically – from an average of about 5.7 kbp to about 30 kbp (see Supplemental Dataset 2 for detailed assembly statistics on each dataset). Even for datasets for which slicing decreased the number reads (Hv5 and Hv10), the n50 increased significantly. The number of reads used by VELVET increased from an average of 77% to 92%; the fraction of known genes that were recovered by the assemblies increased from 81% to 85%. We recognize that the improvement from Table 5 to Table 6 is not just due to the slicing, but also to the increased *k* (from 26 to 32). We have already addressed this point in Table 2, Table 3 and Table 4, where we showed that increasing *k* from 26 to 32 helps the decoding/assembly but the main boost in accuracy and quality is due to slicing.

**Table 6:** Decoding and assembly statistics for the ten datasets using *k*=32 and slicing

| *slicing*<br>no. slices×size | *decoding* | | *assembly* | | | |
| --- | --- | --- | --- | --- | --- | --- |
|  | *reads* (M) | *coverage* (x) | *n50* (bp) | *reads used* (%) | *sum/size* (%) | *observed/expected genes* (%) |
| Hv3 (11×2M) | 1,156.0 | 460.0 | 28,477 | 90.4% | 123.0% | 1,437/1,471 (97.69%) |
| Hv4 (8×2M) | 595.6 | 205.9 | 28,341 | 93.9% | 114.5% | 319/473 (67.44%) |
| Hv5 (4×4M) | 471.0 | 155.5 | 31,038 | 93.6% | 101.0% | 196/226 (86.73%) |
| Hv6 (6×1.5M) | 243.0 | 81.1 | 25,194 | 92.9% | 89.4% | 206/244 (84.43%) |
| Hv7 (15×3M) | 785.0 | 264.0 | 39,742 | 91.1% | 104.0% | 204/228 (89.47%) |
| Hv8 (12×2M) | 669.0 | 223.0 | 34,262 | 91.6% | 102.0% | 187/207 (90.34%) |
| Hv9 (14×1.25M) | 516.0 | 246.0 | 32,634 | 94.3% | 103.2% | 309/361 (85.60%) |
| Hv10 (4×5M) | 695.0 | 499.0 | 42,819 | 90.7% | 121.9% | 20/27 (74.07%) |
| Vu1 (12×1.5M) | 907.0 | 232.0 | 16,388 | 89.7% | 89.7% | 510/612 (83.33%) |
| Vu2 (14×1.5M) | 970.0 | 283.0 | 20,748 | 91.5% | 93.6% | 446/503 (88.67%) |
| *Average* | 700.8 | 265.0 | 29,964 | 92.0% | 104.2% | (84.78%) |

As a final step, we investigated how the depth of sequencing affects BAC assembly quality. To this end, we multiplexed sixteen barley BACs on one lane of the Illumina HiSeq2000, using custom multiplexing adapters. The size of these BACs ranged ≈70 kbp to ≈185 kbp (see Table 7). After demultiplexing the sequenced reads, we obtained 34.4M 92-bases paired-end reads (insert size of 275 bases). We quality-trimmed the reads, then cleaned them of spurious sequencing adaptors; finally reads affected by *E. coli* contamination or BAC vector were discarded. The final number of cleaned reads was 23.1M, with an average length of ≈88 bases. The depth of sequencing for the sixteen BACS ranged from ≈6,600x to ≈27,700x (see Table 7).

**Table 7:** Basic statistics on the read datasets for the 16 barley BACs sequenced individually

| *BAC* | ≈ size (bp) | *set* | *after demultiplexing* | | % *E.coli* | *after demux/cleaning/trimming* | | |
| --- | --- | --- | --- | --- | --- | --- | --- | --- |
|  |  |  | *reads* (M) | *bases* (Mbp) |  | *reads* (M) | *bases* (Mbp) | *coverage* (x) |
| **052L22** | 105,788 | Hv4 | 21.534 | 1,981 | 10.94% | 16.795 | 1,488 | 14,065 |
| **152O10** | 117,543 | Hv3, Hv9 | 18.575 | 1,709 | 16.93% | 13.101 | 1,154 | 9,812 |
| **192B13** | 112,841 | Hv4 | 19.581 | 1,801 | 12.20% | 13.950 | 1,225 | 10,853 |
| **574B01** | 92,859 | Hv3 | 32.952 | 3,032 | 15.27% | 23.414 | 2,059 | 22,172 |
| **630P05** | 110,490 | Hv3 | 16.224 | 1,493 | 15.45% | 11.102 | 971 | 8,792 |
| **727J05** | 131,648 | Hv8 | 22.613 | 2,080 | 14.80% | 15.801 | 1,388 | 10,546 |
| 772L04 | 116,367 | Hv3, Hv10 | 21.837 | 2,009 | 17.44% | 14.407 | 1,255 | 10,784 |
| 773A02 | 185,718 | Hv3, Hv10 | 21.756 | 2,002 | 15.86% | 14.909 | 1,309 | 7,049 |
| 773F12 | 96,385 | Hv3, Hv10 | 20.868 | 1,920 | 14.61% | 14.661 | 1,286 | 13,341 |
| 773H21 | 71,701 | Hv3, Hv10 | 23.679 | 2,178 | 16.38% | 16.770 | 1,473 | 20,540 |
| 773L22 | 92,859 | Hv3, Hv10 | 17.031 | 1,567 | 16.16% | 12.282 | 1,081 | 11,639 |
| 774D07 | 117,543 | Hv3, Hv10 | 13.077 | 1,203 | 20.51% | 8.880 | 778 | 6,621 |
| 774G18 | 90,508 | Hv3, Hv10 | 24.571 | 2,261 | 15.89% | 16.763 | 1,469 | 16,229 |
| 774L04 | 103,437 | Hv3, Hv10 | 22.053 | 2,029 | 17.13% | 15.219 | 1,334 | 12,895 |
| 774O01 | 95,209 | Hv3, Hv10 | 41.579 | 3,822 | 14.76% | 29.877 | 2,642 | 27,754 |
| **789L09** | 84,631 | Hv3 | 37.000 | 3,404 | 15.20% | 25.730 | 2,264 | 26,754 |

Another set of 52 barley BACs was sequenced by the Department of Energy Joint Genome Institute (JGI) using Sanger long reads. All BACs were sequenced and finished using PHRED/PHRAP/CONSED to a targeted depth of 10x. The primary DNA sequences for each of these 52 BACs was assembled in one contig, although two of them were considered partial sequence.

The intersection between the set of 16 BACs sequenced using the Illumina instrument and the set of 52 BACs sequenced using Sanger is a set of seven BACs (highlighted in bold in Table 7), but one of these seven BACs is not full-length (052L22). We used the six full-length Sanger-based BAC assemblies as the “ground truth” to assess the quality of the assemblies from Illumina reads at increasing depth of sequencing. To this end, we generated datasets corresponding to 100x, 250x, 500x, 1,000x, 2,000x, 3,500x, 5,000x, 6,000x, 7,000x and 8,000x depth of sequencing (for each of the six BACs), by sampling uniformly short reads from the high-depth datasets. For each choice of the depth of sequencing, we generated twenty different datasets, for a total of 1,200 datasets. We assembled the reads on each dataset with VELVET v1.2.09 (with hash value *k* = 79 to minimize the probability of false overlaps) and collected statistics for the resulting assemblies. Figure 4 shows the value of n50 (A), the size of the largest contig (B), the percentage of the target BAC not available in the assembly (C), and number of assembly errors (D) for increasing depth of sequencing. Each point in the graph is the average over the twenty datasets, and error bars indicate the standard deviation. In order to compute the number of assembly errors we used the tool developed for the GAGE competition [13]. According to GAGE, the number of assembly errors is defined as the number of locations with insertion/deletions of at least six nucleotides, plus the number of translocations and inversions.

**Figure 4:**
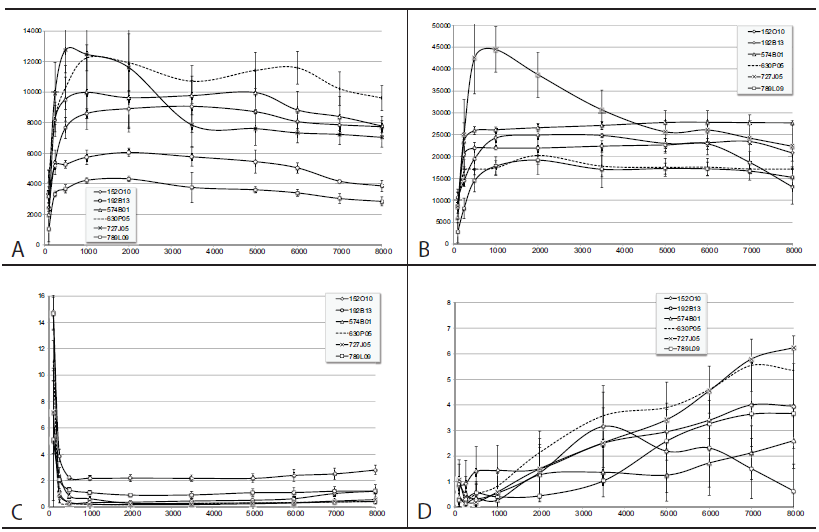
VELVET assembly statistics as a function of the depth of sequencing coverage: (A) N50, (B) longest contig, (C) percentage of the target BAC not covered by the assembly, (D) number of assembly errors; each point is an average over twenty samples of the reads, errors bars indicate standard deviation among the samples

A few observations on Figure 4 are in order. First, note that both the n50 and the size of the longest contig reach a maximum in the 500x-2000x range, depending on the BAC. Also observe that in order to minimize the percentage of BAC missed by the assembly one needs to keep the depth of sequencing below 2,500x (too much depth decreases the coverage of the target). Finally, it is very clear from (D), that as the depth of sequencing increases so do the number of assembly errors (with the exception of one BAC).

We have also investigated whether similar observations could be drawn for other assemblers. In Figure 5, we report the same assembly statistics, namely (A) the value of n50, (B) the size of the largest contig, (C) the percentage of the target BAC not available in the assembly, and (D) number of assembly errors for increasing depth of sequencing for one of the BACs. This time we used three assemblers, namely VELVET, SPADES v3.1.1 [14] and IDBA-UD [15] (statistics for all BACs are available in Supplemental Text, Figures S2 – S5). While there are performance differences among the three assemblers, the common trend is that as the coverage increases, the N50 and the size of the largest contig decreases, while the percentage of the BAC missing and the number of assembly errors increases. Among the three assemblers, SPADES appears to be less affected by high coverage, but still its performance does not improve with higher and higher coverage, as one would expect. We should note that SPADES has a sophisticated error-correction preprocessing step. SPADES was run with hash values *k* = 25, 45, 65 and option-careful (other parameters were default). IDBA-UD was run with hash values *k* = 25, 45, 65 (other parameters were default). The reported assembly is the one chosen by IDBA-UD.

**Figure 5:**
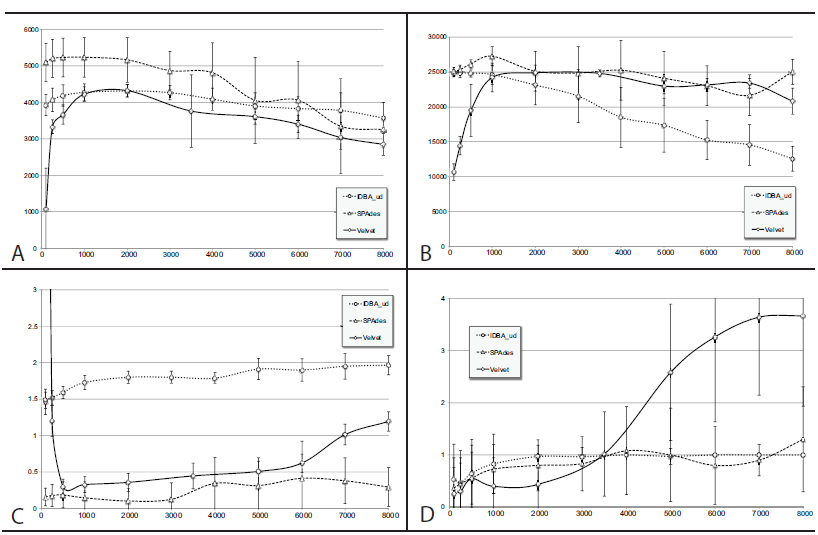
Assembly statistics as a function of the depth of sequencing coverage for BAC 789L09 for three assemblers: VELVET, SPADES and IDBA; (A) N50, (B) longest contig, (C) percentage of the target BAC not covered by the assembly, (D) number of assembly errors; each point is an average over ten subsamples of the reads, errors bars indicate standard deviation among the samples

Finally, we analyzed the performance of IDBA-UD and VELVET on simulated reads. We generated 100bp×2 paired-end reads from the Sanger assembly of BAC 574B01 using the read simulator WGSIM (https://github.com/lh3/wgsim) at 100x, 250x, 500x, 1,000x, 2,000x, 3,500x, 5,000x, 6,000x, 7,000x and 8,000x depth of sequencing. Insert length was 250bp, with a standard deviation of 10bp. For each depth of sequencing, we generated simulated reads at 0%, 0.5%, 1%, and 2% sequencing error rate (substitutions). Insertions and deletions were not allowed.

IDBA-UD was executed with hash values *k* = 25, 45, 65 (other parameters were default). VELVET was run with *k* = 49. We repeated the simulations twenty times for IDBA-UD and ten times for VELVET. In the Supplemental Text (Figure S5–S6) we report the usual assembly statistics, namely N50, largest contig, percentage missing, and number of assembly errors for IDBA-UD and VELVET on these datasets. Observe that with “perfect” reads (0% error rate), ultra-deep coverage does not affect the performance of IDBA-UD and VELVET. With higher and higher sequencing errors, however, similar behaviors to the assembly of real data can be observed: N50 and longest contig rapidly decrease, and missing portions of the BAC and number of mis-assemblies increase.

## Discussion

Since the introduction of DNA sequencing in the seventies, scientists had to come up with clever solutions to deal with the problem of *de novo* genome assembly with limited depth of sequencing. As the cost of sequencing keeps decreasing, one can expect that computational biologists will have to deal with the opposite problem: excessive amount of sequencing data. The Lander-Waterman-Roach theory [16], [17] is a good model to estimate gap and contig lengths when the depth of sequencing is low to moderate, but does not explain why the quality of the assembly starts degrading when the depth is too high. Our conjecture is that in the presence of sequencing errors, the more sequencing data we have, the more likely it is to observe two or more reads with sequencing errors in the same positions, which in turns increases the chance of detecting false overlaps. These false overlaps induce mis-assemblies or become seeds for spurious contigs. This conjecture is supported by the simulation results discussed at the end of the previous section.

In this study, we report on the *de novo* assembly of BAC clones, which are relatively short DNA fragments (100–150 kbp). With current sequencing technology it is very easy to reach depth of sequencing in the range of 1,000x–10,000x and study how the assembly quality changes as the amount of sequencing data increases. Our experiments show that when the depth of sequencing exceeds a threshold the overall quality of the assembly starts degrading (Figure 4). This appears to be a common problem for several *de novo* assemblers (Figure 5). The same behavior is observed for the problem of decoding reads to their source BAC (Figure 2), which is the main focus of this manuscript.

The next question is how to deal with the problem of excessive sequencing depth. For the decoding problem we have presented an effective “divide and conquer” solution: we “slice” the data in subsamples, decode each slice independently, then merge the results. In order to handle conflicts in the BAC assignments (i.e., reads that appear in multiple slices that are decoded to different sets of BACs), we devised a simple set of voting rules. The question that is still open is what to do for the assembly problem: one could assemble slices of the data independently, but it is not clear how to merge the resulting assemblies. There is a growing set of literature on assembly reconciliation/merging (see, e.g., [18], [19], [20]) but it is not obvious how to apply these methods to merge a large number of independent assemblies.

At the time of writing we are working on an iterative assembler that employs a strategy similar to the voting method used here for decoding. In this method, we assemble reads in each independent slice using an off-the-shelf assembler (e.g., SPADES), then identify sufficiently long DNA segments that are shared by the majority of the assemblies. These *conserved segments* are added to the *consensus* assembly (which is initially empty). Before starting the second iteration, any read that maps to conserved segments is removed from the input dataset. The reduced input dataset is sliced again, and the process is repeated until no sufficiently long conserved regions can be found. An alternative approach to deal with ultra-deep coverage would be to attempt to eliminate from the dataset reads that are affected by excessive sequencing errors, but this problem is not easy to solve when a reference genome is not available. In general, we believe that the problem of *de novo* sequence assembly must be revisited from the ground up under the assumption of ultra-deep coverage.

## Author contributions

SL and TJC designed and supervised the project. SL wrote the draft of the manuscript, designed the algorithm for conflict resolution, and collected the experimental results. SHM collected comparative assembly results for the six BACs with high depth of sequencing. SW wrote the scripts to demultiplex and clean/trim the sequencing data, the script that slices the pools, and the script that runs BOWTIE on the decoded reads against the 454-based BAC assemblies. SW suggested the idea of making the size of the pools more “balanced”, which led to the exploration of the slicing size. MA wrote scripts to further clean the reads, assemble the BACs, and BLAST unigenes against BAC assemblies. GC suggested the explanation that too much data can hurt the decoding algorithm, participated in the discussions about slicing, and edited the manuscript.

AD suggested some the analyses used in this manuscript. TJC provided the sequencing data for barley and cowpea, supervised the project, and edited the manuscript. All authors read and approved the final manuscript.

## Description of additional data files and software

Raw reads for barley and cowpea BACs have been deposited in NCBI SRA accession number SRA051780, SRA051535, SRA051768, SRA073696, SRA051739 (barley); SRA052227 and SRA052228 (cowpea).

Python scripts to process slices and resolve the decoding conflicts are available from http://goo.gl/YXgdHT The latter archive also includes a small dataset of reads that can be used to test whether the software pipeline is working on a specific computer architecture. Software HASHFILTER can be downloaded from http://goo.gl/MIyZHs

## Acknowledgments

This work was supported in part by the U. S. National Science Foundation [DBI-1062301], by the USDA National Institute of Food and Agriculture [2009-65300-05645] and by the USAID Feed the Future program [AID-OAA-A-13-00070].

We thank the Department of Energy Joint Genome Institute (Dr. Jane Grimwood, Dr. Jeremy Schmutz) for the use of the reference BAC barley clone data 14090–14118 assembled from Sanger sequencing data. We also thank Prof. Titus Brown (Michigan State U.) for useful comments on the manuscript.

